# Identification of determinants of high-fidelity DNA synthesis in *M. smegmatis* DnaE1 through *in silico* and *in vivo* approaches

**DOI:** 10.1101/2025.04.17.649371

**Authors:** R.C.M. Kuin, G.J.P. van Westen, M.H. Lamers

## Abstract

Drug resistance in *Mycobacterium tuberculosis* presents a major challenge in tuberculosis treatment, highlighting the need to understand the underlying mechanisms. DNA replication plays an important role in the acquisition of drug resistance and the expression of the DNA polymerase DnaE2 during adverse conditions has been associated with increased mutation rates. Here we investigate the functional differences between the high fidelity replicative DNA polymerase DnaE1 and the predicted error-prone DNA polymerase DnaE2, focusing on which amino acid changes affect polymerase fidelity. For this we identify potential fidelity-altering positions using a two-entropies sequence analysis combined with experimental validation to test whether changes of these positions affect the mutation rates. We find that a double mutation in the palm domain of *M. smegmatis* DnaE1: D431S/R432D, increases mutation frequencies both *in vivo* and *in vitro*. The location of these two residues adjacent to the DNA backbone of the template strand suggests that the amino acid change results in a looser grip on the DNA, allowing for the incorporation of incorrect nucleotides. These insights improve our understanding of the mechanisms underlying drug resistance in *M. tuberculosis* and could help in the development of future strategies to combat it.

## INTRODUCTION

*Mycobacterium tuberculosis*, the pathogen responsible for tuberculosis, remains the top infectious killer globally (1, 2). The emergence of resistant forms of tuberculosis and in particular multidrug- and extensively drug-resistant tuberculosis form a growing threat to global health (3, 4). These resistant forms of tuberculosis require lengthy treatment regimens with significant side effects, and consume a substantial amount of the healthcare budget and associated resources in endemic countries (5, 6). Hence, a better understanding of the mechanisms behind drug resistance is needed to be able to develop therapeutic interventions to slow down drug resistance.

Unlike many other bacteria, resistance in *M. tuberculosis* is not acquired via horizontal gene transfer (7, 8). Instead, the driver of drug resistance in *M. tuberculosis* is the acquisition of mutations during DNA replication in genes that encode drug targets or drug-activating enzymes (7, 9). Therefore, targeting DNA replication and more specifically the bacterial DNA polymerase would be an attractive approach for slowing down drug resistance (10, 11).

Mycobacteria contain two copies of a C-family DNA polymerase: the high-fidelity replicative DNA polymerase DnaE1 and DNA polymerase DnaE2, which is expressed under adverse conditions, such as DNA damage or antibiotic exposure (8, 12–14) While DnaE1 ensures accurate DNA replication, the expression of DnaE2 has been associated with increased mutation rates, reflecting its role in adaptive responses (13, 15). Unlike DnaE1, which is functionally organized into the replisome, DnaE2 interacts with other proteins—ImuA’ and ImuB—to form the mutasome, a complex crucial for its error-prone activity (16).

Like other C-family DNA polymerases, DnaE2 contains both a polymerase active site and an exonuclease active site (17, 18). The importance of the polymerase active site is highlighted by the observation that changes in the polymerase active site of DnaE2 reduce mutation rates in *M. tuberculosis* (14). In contrast, the exonuclease domain of DnaE2 is predicted to be inactive due to the absence of a critical metal-binding residue (16). The predicted lack of a functional exonuclease domain in DnaE2 fits well with its association with higher mutation rates as inactivating mutations in the exonuclease domain of DnaE1 lead to increased mutation rates in the bacterium (16, 19, 20). However, whether DnaE2 also incorporates nucleotides with reduced accuracy remains an open question.

So far, no biochemical studies involving DnaE2 have been published, possibly due to the difficulty of working with DnaE2 in isolation (15). To work around these challenges, we employed a two-entropies analysis of 358 DnaE1 and DnaE2 sequences to identify amino acid positions that potentially influence fidelity of DNA synthesis. Next, we created 16 DnaE1 variants by replacing these amino acids with their corresponding residue from DnaE2 and tested these in *M. smegmatis* through dCas9-mediated knockdown of endogenous DnaE1 and rescue by the variant DnaE1. Variants with increased mutation rates were subsequently purified and analysed *in vitro* for fidelity of DNA synthesis.

Our results identify a double mutant in the palm domain of *M. smegmatis* DnaE1: D431S/R432D, which exhibits enhanced mutation frequencies in both phenotypic and biochemical settings. The location of the double mutation adjacent to the backbone of the template strand suggests that the alteration may create a more open active site that is more generous to mis-incorporated nucleotides. We also confirmed the predicted lack of exonuclease activity in DnaE2 by changing key residues in the PHP domain that is responsible for exonuclease activity in DnaE1. Combined, these results indicate that DnaE2 acts as an error-prone polymerase that introduces more errors during catalysis, providing novel insights into the functioning of the *M. smegmatis* mutasome.

## METHODS

### Materials

All chemicals were purchased from Sigma-Aldrich unless stated otherwise. DNA substrates were ordered from IDT.

### Two-entropies analysis of DnaE1 and DnaE2 sequences

Homologous DnaE1 and DnaE2 sequences were obtained using BLAST (21). To mitigate bias, we included one sequence per organism. The taxonomy was restricted to only include Mycobacteria that are known to contain DnaE1 and DnaE2 genes (17, 22). The resulting 358 sequences (179 per family) were aligned using ClustalOmega (23). Subsequently, we used the Shannon Entropy to measure sequence conservation at each amino acid position in the sequence alignment, as given by Formula 1 below. Here, the Shannon Entropy (SE) at position *i* in the multiple sequence alignment is given by *a* that loops over the 20 different amino acids and *f*_*ia*_ is the fraction of residues of type *a* at alignment position *i*. Each gap was treated as a different residue so that the conservation for highly gapped positions is low.

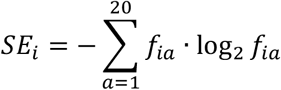

The Shannon Entropy calculations were done for all DnaE1 and DnaE2 sequences combined as well as for DnaE1-only and DnaE2-only sequences separately, using a two-entropies analysis as described (24, 25). The entropy values for DnaE1 and DnaE2 were summed, and values were scaled between 0 and 1. By adding the separate entropy values of DnaE1 and DnaE2, differentially conserved residues can be identified by focusing on residues that have a low protein-specific entropy value and a higher overall entropy value.

### Selection of final set of residues for experimental validation

For making a final selection of differentially conserved residues for experimental validation, we focused on residues with different physicochemical properties between corresponding residues in DnaE1 and DnaE2. For this we use different physicochemical descriptor variables, called Z-scales (26). To measure the variability of Z-scales in the multiple sequence alignment, we calculated the standard deviation per position and scaled these values between 0 and 1.

In addition, to exclude mutations that are predicted to have a negative impact on protein stability, we estimated the changes in free energy upon mutation for all possible DnaE1 to DnaE2 mutations using ICM-Pro version 3.9-3a (Molsoft L.L.C.).(27) The free energy change (ΔΔG) in protein stability was then calculated as the difference between the Gibbs energy (ΔG) of the mutant and the wild type. In a similar way, the predicted change in binding free energy (ΔΔGbind) to the DNA was calculated upon mutation. The final selection was made by visual inspection and selecting residues close the DNA, as these were expected to have biggest effect on fidelity.

### Preparation of dCas9 knockdown plasmids

Knockdown of *M. smegmatis* DnaE1 was achieved by using a catalytically dead Cas9 (dCas9). sgRNA oligo’s were designed using the Pebble design workflow (28). The primer pair was annealed and cloned into the anhydrotetracycline (ATc) inducible vector plJR962 (Addgene: #115162) using restriction cloning as described (28, 29). Briefly, plJR962 was digested with BsmBI-v2 and gel purified. Two complementary oligonucleotides were then annealed and cloned into the digested plasmid backbone. Different oligo pairs were tested, and the one predicted to be the strongest repressor was selected for DnaE1 (Top: 5’-GGGAAGACCGTGGTCCATCAGTTCGAG-3’, Bottom: 5’-AAACCTCGAACTGATGGACCACGGTCT-3’), targeting the PAM sequence corresponding to residues 196–198 of DnaE1. As a control for increased mutagenesis we targeted the EndoMS/NucS enzyme that is responsible for the post-replicative removal of mismatches in mycobacteria (30). A NucS targeting primer pair (Top: 5’-GGGAGGCGAACACACCTGCCACCGGCG-3’, Bottom: 5’-AAACCGCCGGTGGCAGGTGTGTTCGCC-3’) was obtained and cloned as described above. Plasmid construction was performed in *E. coli* DH5α cells, grown in Luria-Bertani (LB) medium at 37 °C. Antibiotics were used at the following concentrations: kanamycin at 50 µg/ml, chloramphenicol at 35 µg/ml, and hygromycin at 200 µg/ml.

### Preparation of CRISPRi-Resistant DnaE1 variant plasmids

The *M. smegmatis* DnaE1 gene (UniProt ID: A0QX55) was cloned in the pINIT vector (Addgene: #46858) using FX-compatible primers and the FX protocol (31). Two silent mutations in codons 196 (AAC → AAT) and 198 (TTC → TTT) were introduced in the sgRNA target sequence of the DnaE1 expression plasmid to make it CRISPRi resistant without changing the amino acid sequence. DnaE1 mutants were generated by using commercial dsDNA Gene Fragments (IDT) and cloning them in the sgRNA-resistant pINIT vector using In Vivo Assembly (32). The Gene Fragments and associated primers were designed using the Python package PyVADesign (33) Mutants were then subcloned in a modified pACE expression vector without GFP tag for acetamide-inducible expression using FX cloning (31). Plasmid construction was performed in *E. coli* DH5α cells, as described above.

### Preparation of *M. smegmatis* cells expressing dCas9 and DnaE1 variants

*M. smegmatis* MC^2^155 were transformed with the dCas9 plasmid and DnaE1 plasmid in two steps. Cells were made competent as described (34), transformed with 1 µg dCas9 plasmid by electroporation and plated on 7H10 agar plates. Single colonies were selected and made competent a second time and subsequently transformed with the DnaE1 variant plasmid. The resultant colonies were used for whole cell analysis of DnaE1 variants. *M. smegmatis* MC^2^155 was grown aerobically at 37 °C in Middlebrook 7H9 medium containing 0.2% (v/v) glycerol and supplemented with 10% (v/v) albumin-dextrose-catalase (ADC) growth supplement and 0.005% (v/v) Tween-80. Solid media consisted of 7H10 agar containing 0.2% (v/v) glycerol, 10% (v/v) ADC. Antibiotics were used at the following concentrations: kanamycin at 50 µg/ml, hygromycin at 100 µg/ml and rifampicin at 200 µg/ml.

### Whole cell analysis of DnaE1 mutants

*M. smegmatis* MC^2^155 cells transformed with dCas9 and a DnaE1 variant were analysed for both growth defects and changes in mutagenesis rates. To measure the effect of the DnaE1 variant on bacterial growth, cells were recovered for three hours after transformation at 37 °C while shaking at 200 RPM and the culture was split into four parts. Each part was plated on solid media containing 0.4% (w/v) acetamide for DnaE1 expression and/or 100 ng/ml anhydrotetracycline for dCas9 expression. Plates were imaged four days after transformation and the experiment was repeated independently three times.

To measure the effect of the DnaE1 variants on mutagenesis, we counted the number of rifampicin-resistant colonies as an indirect measure of the mutagenesis rate, following the method previously reported (13, 35). For this, cells expressing dCas9 and a DnaE1 variant were grown in a liquid culture to an OD_600_ of 0.4-0.5. Next, 5 ml was plated onto 7H10 solid media containing 200 μg/ml rifampicin. As a control, a dilution series was spotted on solid media without rifampicin to estimate the colony forming unit (CFU). Colonies were counted after four days and mutation frequencies were calculated by dividing the number of rifampicin resistant colonies by the average CFU/ml of the dilution series. Each experiment was performed in triplicate and repeated in three independent measurements.

### Protein purification

*M. smegmatis* DnaE1 WT and mutants, containing an N-terminal His-tag, were expressed in *M. smegmatis* MC^2^155 cells. Cells were grown in 1xYT media at 37 °C while shaking at 200 RPM and protein production was induced at OD_600_ ∼ 0.7 with 0.4% (w/v) acetamide for 6 hours at 30 °C. Cells were lysed by sonication and proteins were purified using a Histrap column (Cytiva) using 25-500 mM Imidazole, 50 mM HEPES pH 7.5, 500 mM NaCl and 1 mM DTT, followed by an heparin column (Cytiva) using 50 mM HEPES pH 7.5, 0.1-1.0 M NaCl and 1 mM DTT. The obtained proteins were found to be at a concentration of 20-30 μM, flash frozen in liquid nitrogen and stored at -70^°^C until use.

### Gel-based DNA polymerase assay

The polymerase assay was performed in 50 mM HEPES pH 7.5, 50 mM potassium glutamate, 6 mg/ml BSA, 2 mM DTT and 5 mM MgCl_2_. Reactions were performed at 20 °C with 50 nM purified protein and 50 nM DNA substrate (For DNA sequences see Supplementary Table S1). Primer extensions were done for 30 minutes in the presence of 100 μM dNTPs (each) and increasing MnCl_2_ concentrations from 0.3 to 10 mM. Reactions were stopped in 50 mM EDTA pH 7.4, separated on a 14% native acrylamide gel, stained with SYBR Safe and imaged with a Bio-Rad Gel Doc XR+ system. Three independent experiments were performed.

## RESULTS

### Design of DnaE1 variants for altered fidelity based on two-entropies analysis

To identify positions that could affect the fidelity of DnaE1, we used a multiple sequence alignment of 358 DnaE1 and DnaE2 sequences. To identify positions that are conserved within each polymerase family but are divergent between them, a two-entropies analysis was applied using the Shannon Entropy (SE) as measure for sequence conservation (24, 25). Shannon Entropy calculations were performed for DnaE1-only and DnaE2-only sequences as well as for all sequences combined (Figure 1A). By summing up the separate Shannon Entropy values of DnaE1 and DnaE2 (SE DnaE1 + SE DnaE2) and comparing these to the overall Shannon entropy values SE (DnaE1+DnaE2), differentially conserved residues can be identified by focusing on residues that have a low protein-family specific entropy value and a high overall entropy value. These differentially conserved residues may be crucial for maintaining the specific enzymatic functions of each polymerase, such as polymerase fidelity. The results of the two-entropies analysis, presented in Figure 1B, show a linear plot, which can be attributed to the limited variability between DnaE1 and DnaE2 sequences, that share an average sequence identity of 55%. To make the two-entropies analysis plot easier to interpret, it has been separated into quadrants with the lines crossing at the point where we can discriminate between overall-conserved and subfamily-conserved residues.

**Figure 1.**
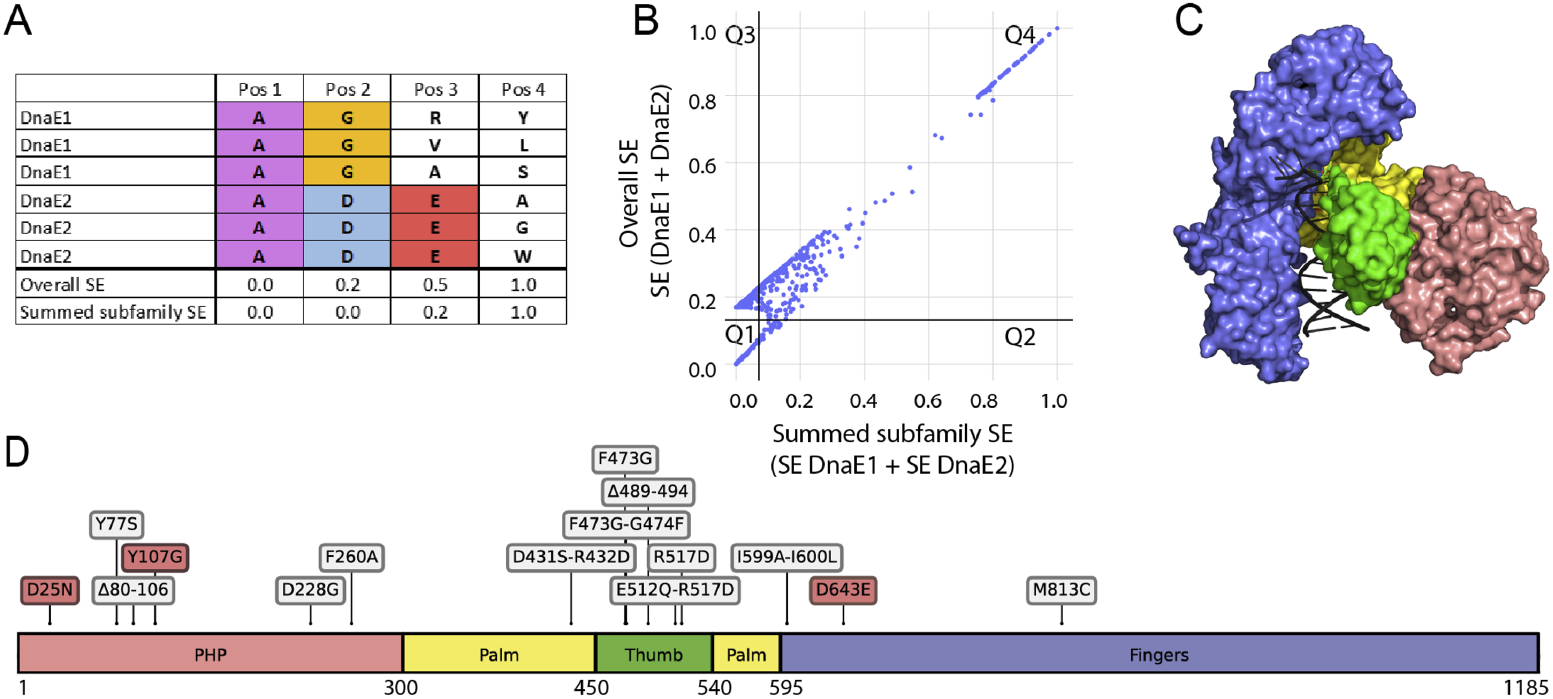
Two-entropies analysis and selection of DnaE1 variants for experimental validation. (**A**) Example positions of a Multiple Sequence Alignment for illustrating the types of positions that were identified in the two-entropies analysis. Residues conserved in both families receive a low score in overall Shannon Entropy (SE) and in the summed subfamily SE (Pos 1). Residues differently conserved per family receive a higher overall SE but a low score for the summed subfamily SE (Pos 2). Residues conserved in only one family are marked by a higher overall SE and slightly elevated summed subfamily SE (Pos 3). Finally, residues that are not conserved in either family are marked by a high overall SE and high summed subfamily SE (Pos 4). (**B**) 2D plot with summed subfamily SE shown on the horizontal axis, and overall SE shown on the vertical axis (scaled between 0 and 1). Each dot represents a position in the Multiple Sequence Alignment of 358 homologous DnaE1 and DnaE2 sequences from Mycobacteria. Two lines (x=0.07 and y=0.13) mark the separation of the residues into four quadrants, based on visual inspection. Example positions Pos 1-4 from (**A**) would approximately correspond to Q1, Q3, Q3, and Q4, respectively. (**C**) Domains (Palm, Fingers, Thumb, PHP) of DnaE1 mapped onto the structure (PDB ID: 7PU7). (**D**) Distribution of the 16 selected DnaE1 variants across the protein sequence, based on the two-entropies analysis. Control variants for increased mutagenesis are shown in red and variants selected based on the two-entropies analysis are shown in grey. The figure was created with DNAFeaturesViewer (47).

Residues in quadrant 1 (Q1, lower left corner) are globally conserved between both polymerase families, such as the *M. smegmatis* polymerase active site residues 424, 426 and 590 that are essential for catalysis in both DnaE1 and DnaE2 (14, 17). In contrast, residues in Q4 (upper right corner), are those that are not conserved in any of the two families. Curiously, Q4 also contains residues located in loops that are present in the DnaE1-family, but not in DnaE2-family polymerases. One of these loops (residues 80-106) is located in the PHP domain (Figure 2, and Figure 1C-D for domain definition). The other loop (residues 489-495) is located in the thumb domain and inserts itself into the major groove of the DNA (12) (Figure 2). To investigate whether these loops affect polymerase fidelity, two DnaE1 variants were designed, each with one of the loops removed.

**Figure 2.**
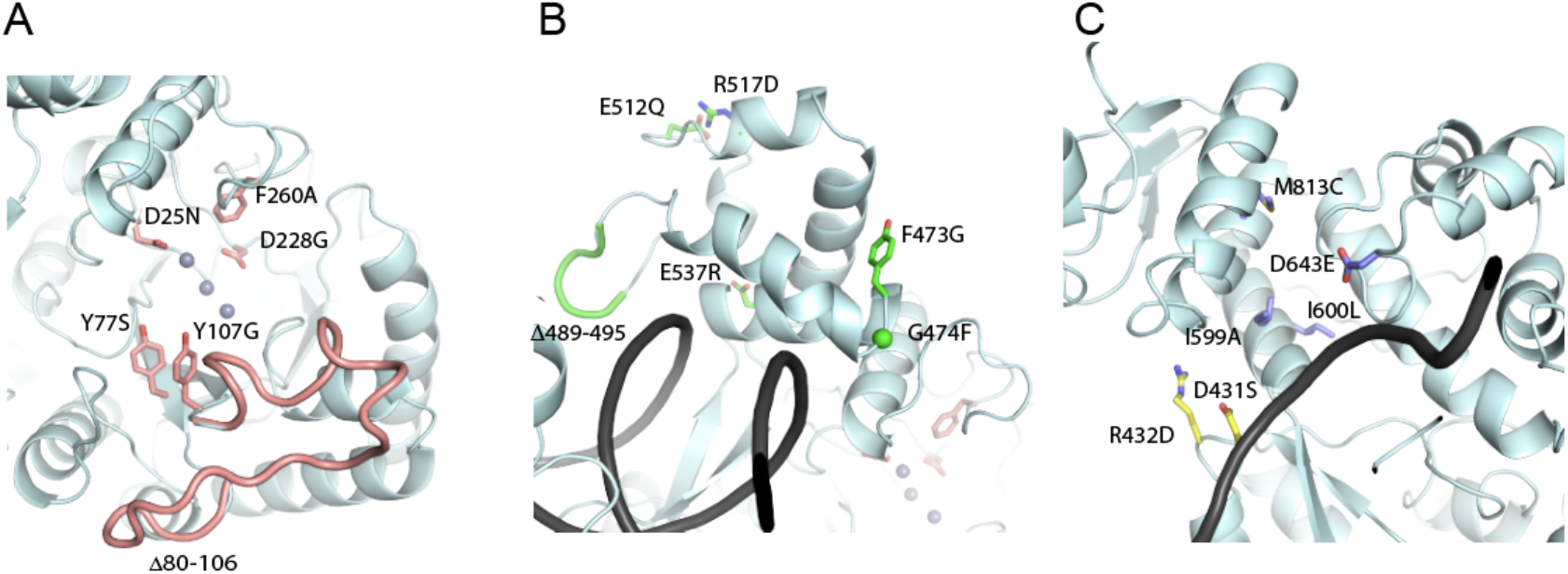
Mutated residues in DnaE1. (**A**) residues in the PHP domain, shown as pink sticks (**B**) residues in the thumb domain, shown as green sticks (**C**) residues in the palm and fingers domain, shown as yellow and purple sticks, respectively.

Quadrant 3 (Q3) shows residues that are conserved in either DnaE1 family or DnaE2 family, but not in both and may therefore play a role in the distinct enzymatic behaviour of the two proteins. It is important to note that differences between DnaE1 and DnaE2 are not only limited to fidelity of DNA synthesis but could also include residues that are required to interact with partner proteins of the replisome or mutasome. Therefore, to maximize the chances of selecting alterations that affect fidelity, we focused on a subset of residues near the DNA binding region. To further enhance the likelihood that alterations impact DNA synthesis, we focused on residues with different physicochemical properties between corresponding residues (Supplementary Fig. S1). Finally, to lower the chances of selecting mutations that could destabilize the protein, the predicted effect on protein stability upon mutation was calculated using the program ICM-Pro (see Methods). Finally, double mutations were designed when two differentially conserved residues were in close proximity to each other in the protein structure to take into account their potential compensatory effect. Based on these criteria, a total of 11 positions were selected for experimental validation (Figure 1C, Supplementary Fig. S1).

To evaluate the effect of the selected positions on polymerase fidelity, variants of DnaE1 were created in *M. smegmatis* by replacing the DnaE1 residue with the corresponding DnaE2 residue. The complete selection of DnaE1 variants contains 11 amino acids variants and two loop deletions based on the two-entropies analysis, and three control mutations based on previous reported effects on DnaE1 (Table 1, Figure 2). Two of the control mutations, D25N and Y107G, are in the PHP domain and were selected based on their inability to remove mis-incorporated nucleotides from a DNA substrate (36). A third control mutation in the fingers domain, D643E, was selected due to the mutator phenotype associated with this mutation in the corresponding residue of *E. coli* Pol IIIα (37). In addition, as positive controls for increased mutagenesis we included a plasmid expressing *M. smegmatis* DnaE2, as well as a dCas9 knockdown of NucS, the key mycobacterial mismatch repair enzyme (30).

**Table 1.** Variants of *M. smegmatis* DnaE1 used in this study.

|  | Mtb<br>residue | Domain | cell growth |  | Mutation frequency |  |
| --- | --- | --- | --- | --- | --- | --- |
|  |  |  | - WT<br>DnaE1 | + WT<br>DnaE1 | - WT<br>DnaE1 | + WT<br>DnaE1 |
| **DnaE1 WT (control)** |  |  | = | = | n.d. | 1.0 |
| **DnaE2 (control)** |  |  | ∅ | = | n.d. | 2.8 ± 0.7 |
| **ΔNucS (control)** |  |  | = | = | n.d. | 64.5 ± 23.7 |
| **D25N (control)** | 23 | PHP | ∅ | < | n.d. | 3.4 ± 2.1 |
| **Y77S** | 75 | PHP | ∅ | < | n.d. | 0.6 ± 0.3 |
| **Δ80-106** | 78-104 | PHP | ∅ | < | n.d. | 0.6 ± 0.3 |
| **Y107G (control)** | 105 | PHP | = | = | 22.1 ± 5.6 | n.d. |
| **D228G** | 226 | PHP | ∅ | < | n.d. | 0.5 ± 0.3 |
| **F260A** | 258 | PHP | ∅ | < | n.d. | 5.9 ± 2.7 |
| **F473G** | 470 | Thumb | ∅ | < | n.d. | 1.9 ± 0.9 |
| **F473G-G474F** | 470-471 | Thumb | ∅ | < | n.d. | 1.2 ± 0.9 |
| **Δ489-495** | 486-492 | Thumb | ∅ | < | n.d. | n.d. |
| **E512Q-R517D** | 509-514 | Thumb | ∅ | < | n.d. | 0.2 ± 0.1 |
| **R517D** | 514 | Thumb | ∅ | < | n.d. | 1.1 ± 0.3 |
| **E537R** | 534 | Thumb | ∅ | = | n.d. | n.d. |
| **D431S-R432D** | 428-429 | Palm | = | = | 7.0 ± 1.9 | n.d. |
| **I599A-I600L** | 596-597 | Fingers | < | = | 0.5 ± 0.1 | n.d. |
| **D643E (control)** | 640 | Fingers | < | = | 0.9 ± 0.8 | n.d. |
| **M813C** | 810 | Fingers | = | = | 0.9 ± 0.5 | n.d. |

Effect on cell growth upon knockdown of endogenous dnaE1 and rescue by variants, measured across three biological replicates. ∅ indicates that there was no growth, < indicates that there was less growth compared to the WT and = indicates that there was equal growth as compared to the WT, four days after transformation. Effect of the DnaE1 variant on mutation frequencies, determined using a rifampicin resistance assay, reported as fold increase (n.d. = not determined) ± the standard deviation across three biological replicates.

### Phenotypic screen using dCas9-mediated knockdown of DnaE1 reveals little tolerance for variation in DnaE1

The DnaE1 variants selected above were tested in *M. smegmatis* through dCas9-mediated knockdown of endogenous DnaE1 and rescue by the variant DnaE1. For ease of use, we used the non-pathogenic, fast dividing *M. smegmatis*, that shows high genetic resemblance to *M. tuberculosis* (Supplementary Fig. S4/S5) and conservation of DNA replication and mutagenesis mechanisms (38, 39). In this setup, knockdown of DnaE1 results in no growth, consistent with the essential role of DnaE1 (40), which can be rescued by plasmid-based expression of dCas9-resistant DnaE1 (Supplementary Fig. S4). Next, we evaluated the effect of the DnaE1 variants on cell growth by growing them in presence and absence of the genomic DnaE1 (Table 1). Surprisingly, the majority of DnaE1 variants could not grow in absence of the genomic DnaE1, indicating that DnaE1 is intolerant to mutations across all domains (PHP, thumb, palm, fingers, see Figure 1C-D). In addition, most DnaE1 variants also showed reduced growth when genomic DnaE1 was present, indicating a dominant negative effect. This suggests that the presence of the mutant polymerase disrupts the DNA synthesis by competing with the wild type DnaE1 for a place in the replisome. We also tested if DnaE2 could replace DnaE1, which resulted in no growth. In contrast, expression of DnaE2 in the presence of WT DnaE1 did not show any impact on growth, unlike many of the DnaE1 variants. This not only suggests that DnaE2 cannot replace DnaE1, but that it also cannot interact with the replisome as it does not exert a dominant negative effect. Instead, DnaE2 has evolved to interact with the mutasome (14).

### The double mutant D431S/R432D shows enhanced mutation frequencies

To measure the effect of the DnaE1 variants on fidelity of DNA synthesis, we used the appearance of rifampicin resistant colonies as a measure for increased mutagenesis (13, 41) The setup of this assay was confirmed by measuring the mutation frequency of a dCas9 knockdown of NucS, the key enzyme of the non-canonical DNA mismatch repair pathway in mycobacteria (30). Knockdown of NucS results in a ∼64 fold increase in mutation frequency (Table 1). Also, the expression of DnaE2 in presence of the genomic DnaE1 increases mutagenesis, but only by 3-fold. The limited impact on mutagenesis upon expression of DnaE2 may be explained by the lack of the other mutasome components, ImuA’ and ImuB, as it was reported that deletion of any of the mutasome components eliminated mutagenesis in *M. smegmatis* (14, 42).

Next, we tested the effect of the DnaE1 variants on mutagenesis. In the PHP domain, only one of the variants could be expressed in the absence of the wild type gene. This control variant, Y107G, shows a 22-fold increase in the number of rifampicin resistant colonies. This is consistent with previous work that showed that this mutant lacks exonuclease activity (36). As the remaining variants in the PHP domain could not rescue the wild type gene, we measured their impact on mutagenesis in the presence of the endogenous DnaE1 (Table 1). We find that co-expression of the D25N and F260A variants with wild type DnaE1 led to a 3.4-fold and 5.9-fold increase in mutagenesis (Table 1), respectively. The smaller impact on mutagenesis compared to Y107G may be explained by the presence of the wild type DnaE1 that attenuates their impact on mutagenesis. Other PHP variants (Y77S, D228G, and Δ80-106) did not increase mutagenesis. The lack of effect for D228G may be due to protein instability (36) and for Y77S and Δ80-106 it could be either be that the mutation does not affect exonuclease activity, or that they result in unstable protein too. Thumb domain variants, expressed only with wild type DnaE1, negatively affected bacterial growth but did not increase mutagenesis, suggesting that their mutation effects or compromised polymerase activity, leading to e.g. stalled DNA synthesis. Variants in the fingers domain, expressed without the wild type protein, showed no impact on mutagenesis, indicating no effect on polymerase fidelity. In contrast, the variant in the palm domain: D431S/R432D, showed a 7-fold increase in mutagenesis, without having an impact on bacterial growth.

Hence, from the 16 variants tested, only two variants, Y107G and D431S/R432D, showed increased mutagenesis without impacting the growth of the bacterium. For further analysis these two variants were purified and their activity was assessed *in vitro*. To do so, we monitored DNA polymerase activity using four specially designed DNA substrates. In each of these, the template strand consists of an 18-nucleotide, single-stranded 3’ overhang containing only three of the four nucleotides with a single fourth nucleotide placed at the centre (Figure 3). Next, by providing only three of the complementary nucleotides, the polymerase will stall on the single nucleotide, unless it is able to insert and extend from a mismatch. To enhance the number of mismatches, we added increasing amounts of manganese, which is known to promote the error-rate of the polymerase (43), thus increasing the likelihood that a mismatch is made.

**Figure 3.**
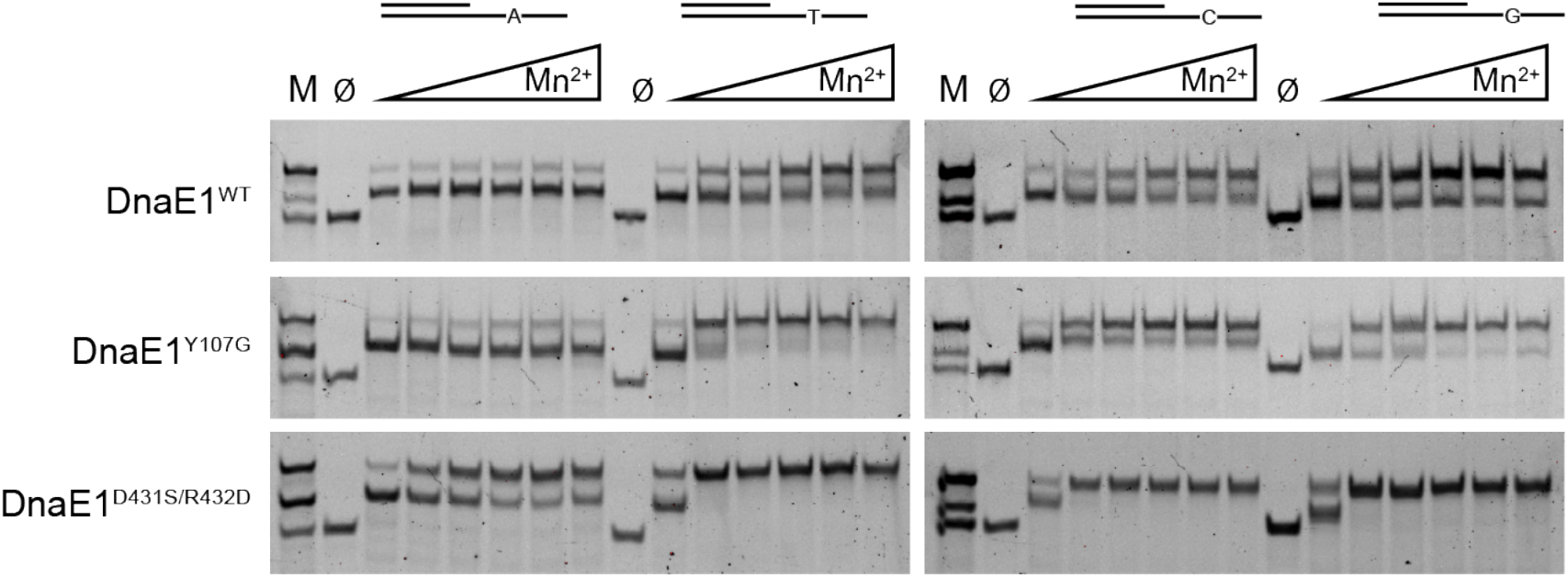
Polymerase assay of wild type and variant *M. smegmatis* DnaE1. Four different substrates were used, each with a unique nucleotide in the single stranded region of the template DNA. Extension reactions were performed with only three of the four nucleotides, thus requiring the polymerase to create a mismatch for complete extension of the primer strand. Increasing amounts of manganese (Mn^2+^: from 0.3 to 10 mM) were added to enhance the error rate of the polymerase. Marker lane (M) show the start substrate, partially extended substrate and fully extended substrate. ∅ marks the lane in which all nucleotides were omitted from the reaction.

For the wild type protein, we find that in the absence of manganese, the polymerase rarely makes it past the middle of the single stranded overhang where the fourth nucleotide is positioned. Addition of manganese increases the amount of fully extended product, although much of the stalled product remains. In the Y107G variant that lacks exonuclease activity (36) we find that the amount of fully extended product is increased compared to the wild type protein. The amount of fully extended product is almost complete in the D431S/R432D variant indicating that this double mutation in the palm domain strongly enhances the propensity of the polymerase to incorporate and extend a mismatch.

Curiously, addition of manganese shows no effect on the substrate with the single A in the single stranded overhang. This is also the case for the Y107G variant. In contrast, the D431S-R432D variant shows a strong increase of the full-length product. This could mean that high fidelity DNA synthesis opposite an A is controlled by insertion fidelity, while insertion opposite the other nucleotides (C, G, T) is more controlled by extension fidelity.

### The location of the D431S/R432D suggests a more open DNA binding groove

As shown above, mutations in the PHP domain result in an increased mutagenesis, which is consistent with various observations that inactivation of the proofreading activity increases mutation rates (16, 36, 44). The location of the mutation-inducing variant D431S/R432D on the other hand is > 40 Å away from the variants in the PHP domain (Figure 4A), suggesting it acts in another way. The two residues are located adjacent to the sugar-phosphate backbone of the template strand, two base pairs away from the insertion site of the incoming nucleotide (Figure 4B). They are located immediately after the strand that holds two of the catalytic residues of the polymerase active site: D424 and D426 (Figure 4B). Interestingly, D431 and R432 are close to, but do not contact the DNA backbone. Instead, they form hydrogen bonds with two resides in the fingers domain: T598 and E562 (Figure 4C). Doing so, the hydrogen bonds stabilize the position of D431 and R432 so that together with R595 they create a groove that follows the DNA backbone (Figure 4D). The increased mutation rate observed for the D431S/R432D variant could be the result of a wider DNA binding groove that enables the DNA to adapt to wrongly incorporated nucleotides. Alternatively, or in addition, the lack of hydrogen bonds in the D431S/R432D variant may allow for increased movement of the catalytic residues D424 and D426, located immediately upstream of the double mutation, that thereby become more generous to mis-incorporated nucleotides.

**Figure 4.**
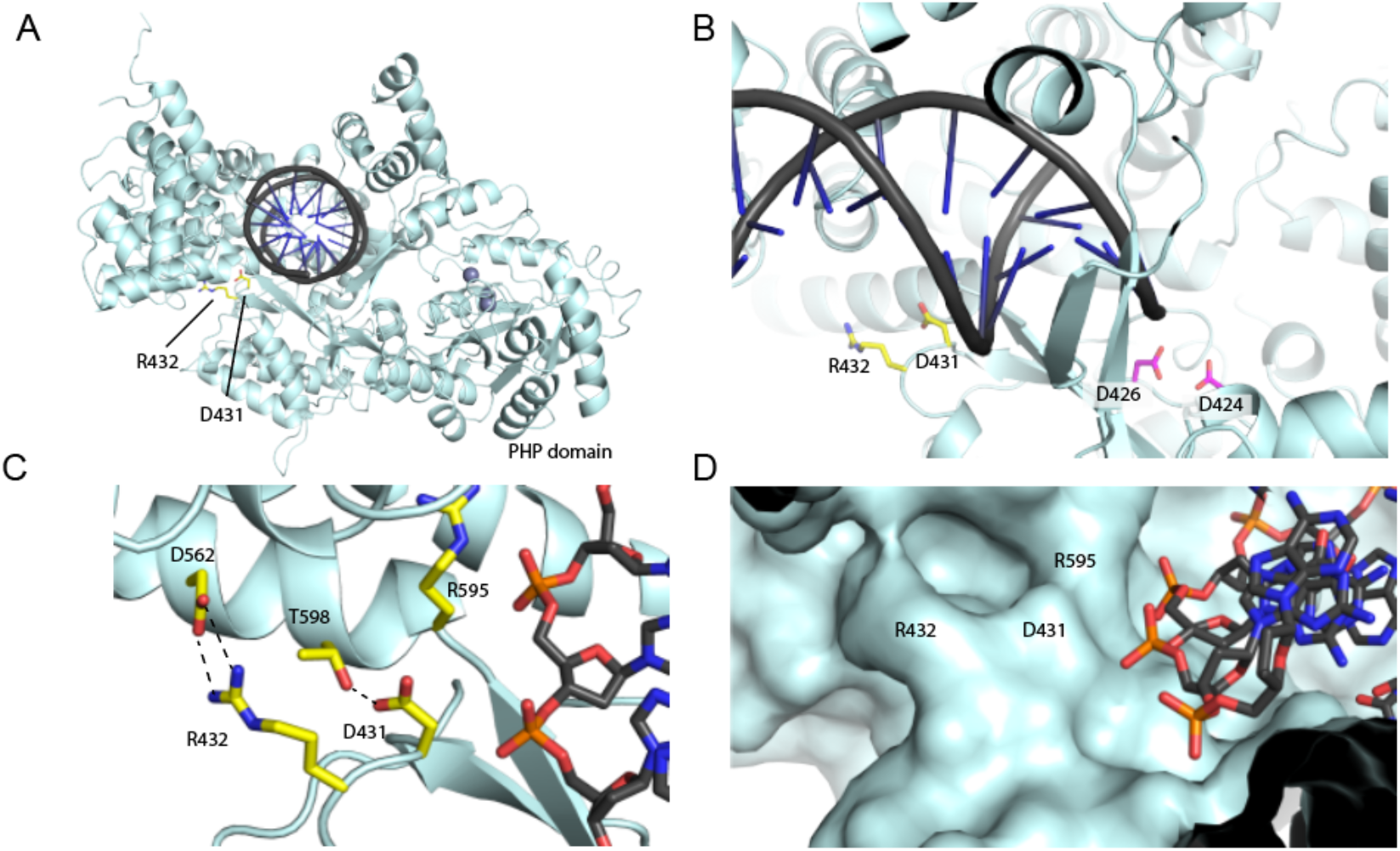
Analysis of DnaE1 D431S/R432D. (**A**) Location of the D431 and R432 in the Palm domain of DnaE1. (**B**) Close up of the position of the two residues adjacent to the template strand and immediately downstream of the catalytic residues D424 and D426 (**C**) Hydrogen bonding between D431 and R432 to two residues in the Fingers domain; T598 and D562. (**D**) Surface representation of the Palm domain, showing a groove for the DNA template strand. Location of D431, R432 and R595 is shown.

## DISCUSSION

Tuberculosis poses a significant global health risk, a situation worsened by the frequent emergence of drug-resistant strains. Resistance is often associated with *de novo* point mutations in the DNA that are introduced during DNA replication (7). To better understand these mechanisms, it is crucial to explore factors that affect mutation rates.

The expression of DnaE2 during adverse conditions has been associated with increased mutation rates and drug resistance in mycobacteria (13). However, due to the challenges of working with DnaE2 in isolation, biochemical studies involving DnaE2 have not yet been published. To work around these challenges, we employed a computational two-entropies analysis of homologous DnaE1 and DnaE2 sequences from mycobacteria to identify positions in the polymerases that could affect the polymerase fidelity. Next, we tested variants of DnaE1, harboring one or more mutations that make it more similar to DnaE2, in *M. smegmatis* through Cas9 knockdown of endogenous DnaE1 and rescue by the variant DnaE1. Variants with increased mutation rates were subsequently purified and analyzed in vitro for fidelity of DNA synthesis.

Polymerase fidelity is determined by two processes: nucleotide insertion fidelity and extension fidelity. The insertion fidelity selects the correct nucleotide opposite the template base. When the polymerase does incorporate an incorrect nucleotide, the distortion of the 3’ of the primer strand slows down or even prevents the incorporation of the next nucleotide, resulting in extension fidelity. Therefore, all high-fidelity DNA polymerases contain an exonuclease, as mis-incorporated nucleotides become roadblocks to DNA synthesis. As a result, inactivation of the exonuclease domain of replicative DNA polymerases leads to severe growth defects (44–46). In contrast, low fidelity Y family DNA polymerases such as the bacterial DNA Pol IV and Pol V, and the eukaryotic DNA polymerases Pol η, Pol κ, and Pol ι do not contain exonuclease domains, as their more open active site allows for both insertion and extension of mis-incorporated nucleotides. It was previously reported that DnaE2 lacks a residue in the PHP domain at the equivalent position of D228 in *M. smegmatis* DnaE1 (or D226 in *M. tuberculosis* DnaE1) that is required for exonuclease activity (16, 36, 44). Therefore, given that a high-fidelity DNA polymerase without exonuclease activity leads to stalled DNA polymerase, it follows that DnaE2 must have a lower insertion and extension fidelity in order to function as a true error-prone DNA polymerase. However, due to the difficulty with working with purified DnaE2, this has not been validated. A pairwise comparison of DnaE1 and DnaE2 sequences shows only 25% sequence identity, and 42% sequence similarity (Supplementary Fig. S3), making the identification of residues that may contribute to high-fidelity DNA synthesis impossible. Here we used a two-entropies analysis of 358 homologous DnaE1 and DnaE2 sequences from mycobacteria to identify 13 variants of DnaE1 that may contribute to DNA polymerase fidelity. Subsequent phenotypic and biochemical analysis revealed two residues in the palm domain, D431 and R432, that lower the fidelity of the polymerase when mutated to their corresponding DnaE2 residue, with no noticeable effect on cell growth. We furthermore show that DnaE1: D431S/R432D also incorporates more errors across different bases in a primer extension assay. These findings strongly suggest that that the insertion fidelity of DnaE2 is lower than DnaE1, providing evidence that DnaE2 is an error-prone polymerase. Due to the high sequence identity of 86% between *M. tuberculosis* DnaE1 and *M. smegmatis* DnaE1, as well as the strong conservation observed in the DnaE2 alignment (Supplementary Fig. S3/S5), we believe that the functional insights gained from *M. smegmatis* will be highly relevant to *M. tuberculosis*. With DnaE2 being a key enzyme in the mycobacterial mutasome that has been linked to the rise of drug resistance in *M. tuberculosis*, our work provides new insights into the mechanism of drug resistant in the world deadliest infectious pathogen.

## ACKNOWLEDGEMENTS

We thank Sophia Gessner and Digby Warner for advice on rifampicin resistance assay and extraction of genomic DNA. We thank Adam Fountain for suggestion of using eBlocks for rapid creation of gene variants and we thank Robbert Q. Kim for his assistance in the creation of mutant plasmids. The authors also thank Roelof van der Kleij for his help using the university IT infrastructure. (47, 48). This work has been supported by the Medical Delta Program “AI for Computational Life Sciences”.

## DATA AVAILABILITY

The data underlying this article will be shared on reasonable request to the corresponding author.

## REFERENCES

1. World Health Organization (2023) Global Tuberculosis Report 2023.

2. Mancuso, G., Midiri, A., Gaetano, S.D., Ponzo, E. and Biondo, C. (2023) Tackling Drug-Resistant Tuberculosis: New Challenges from the Old Pathogen Mycobacterium tuberculosis. Microorganisms, 11, 2277.

3. Maitre, T., Aubry, A., Jarlier, V., Robert, J., Veziris, N., Bernard, C., Sougakoff, W., Brossier, F., Cambau, E., Mougari, F., et al. (2017) Multidrug and extensively drug-resistant tuberculosis. Médecine et Maladies Infectieuses, 47, 3–10.

4. Dheda, K., Mirzayev, F., Cirillo, D.M., Udwadia, Z., Dooley, K.E., Chang, K.-C., Omar, S.V., Reuter, A., Perumal, T., Horsburgh, C.R., et al. (2024) Multidrug-resistant tuberculosis. Nat Rev Dis Primers, 10, 1–27.

5. Liebenberg, D., Gordhan, B.G. and Kana, B.D. (2022) Drug resistant tuberculosis: Implications for transmission, diagnosis, and disease management. Front Cell Infect Microbiol, 12, 943545.

6. Akalu, T.Y., Clements, A.C.A., Wolde, H.F. and Alene, K.A. (2023) Economic burden of multidrug-resistant tuberculosis on patients and households: a global systematic review and meta-analysis. Sci Rep, 13, 22361.

7. Dookie, N., Rambaran, S., Padayatchi, N., Mahomed, S. and Naidoo, K. (2018) Evolution of drug resistance in Mycobacterium tuberculosis: a review on the molecular determinants of resistance and implications for personalized care. Journal of Antimicrobial Chemotherapy, 73, 1138–1151.

8. Salini, S., Bhat, S.G., Naz, S., Natesh, R., Kumar, R.A., Nandicoori, V.K. and Kurthkoti, K. (2022) The Error-Prone Polymerase DnaE2 Mediates the Evolution of Antibiotic Resistance in Persister Mycobacterial Cells. Antimicrobial Agents and Chemotherapy, 66, e01773–21.

9. Pérez-Marinez, D.E. and Zenteno-Cuevas, R. (2024) SNPs in genes related to the repair of damage to DNA in clinical isolates of M. tuberculosis: A transversal and longitudinal approach. PLoS One, 19, e0295464.

10. Santos, J.A. and Lamers, M.H. (2020) Novel Antibiotics Targeting Bacterial Replicative DNA Polymerases. An4bio4cs (Basel), 9, 776.

11. Bosch, B., DeJesus, M.A., Poulton, N.C., Zhang, W., Engelhart, C.A., Zaveri, A., Lavaleje, S., Ruecker, N., Trujillo, C., Wallach, J.B., et al. (2021) Genome-wide gene expression tuning reveals diverse vulnerabilities of M. tuberculosis. Cell, 184, 4579-4592.e24.

12. Ditse, Z., Lamers, M.H. and Warner, D.F. (2017) DNA Replication in Mycobacterium tuberculosis. Microbiol Spectr, 5.

13. Boshoff, H.I.M., Reed, M.B., Barry, C.E. and Mizrahi, V. (2003) DnaE2 Polymerase Contributes to In Vivo Survival and the Emergence of Drug Resistance in Mycobacterium tuberculosis. Cell, 113, 183–193.

14. Warner, D.F., Ndwandwe, D.E., Abrahams, G.L., Kana, B.D., Machowski, E.E., Venclovas, Č. and Mizrahi, V. (2010) Essential roles for imuA’- and imuB-encoded accessory factors in DnaE2-dependent mutagenesis in Mycobacterium tuberculosis. Proceedings of the National Academy of Sciences, 107, 13093–13098.

15. Gessner, S., Martin, Z.A.-M., Reiche, M.A., Santos, J.A., Dinkele, R., Ramudzuli, A., Dhar, N., de Wet, T.J., Anoosheh, S., Lang, D.M., et al. (2023) Investigating the composition and recruitment of the mycobacterial ImuA'–ImuB–DnaE2 mutasome. eLife, 12, e75628.

16. Nasir, N. and Kisker, C. (2019) Mechanistic insights into the enzymatic activity and inhibition of the replicative polymerase exonuclease domain from Mycobacterium tuberculosis. DNA Repair (Amst), 74, 17–25.

17. Timinskas, K., Balvočiūtė, M., Timinskas, A. and Venclovas, Č. (2014) Comprehensive analysis of DNA polymerase III α subunits and their homologs in bacterial genomes. Nucleic Acids Res, 42, 1393–1413.

18. Oakley, A.J. (2019) A structural view of bacterial DNA replication. Protein Science, 28, 990–1004.

19. Sekurova, O.N., Sun, Y.-Q., Zehl, M., Rückert, C., Stich, A., Busche, T., Kalinowski, J. and Zotchev, S.B. (2021) Coupling of the engineered DNA “mutator” to a biosensor as a new paradigm for activation of silent biosynthetic gene clusters in Streptomyces. Nucleic Acids Research, 49, 8396–8405.

20. Cai, N., Chen, J., Gao, N., Ni, X., Lei, Y., Pu, W., Wang, L., Che, B., Fan, L., Zhou, W., et al. (2023) Engineering of the DNA replication and repair machinery to develop binary mutators for rapid genome evolution of Corynebacterium glutamicum. Nucleic Acids Research, 10.1093/nar/gkad602.

21. Altschul, S.F., Gish, W., Miller, W., Myers, E.W. and Lipman, D.J. (1990) Basic local alignment search tool. J Mol Biol, 215, 403–410.

22. Timinskas, K. and Venclovas, Č. (2019) New insights into the structures and interactions of bacterial Y-family DNA polymerases. Nucleic Acids Research, 47, 4393–4405.

23. Sievers, F. and Higgins, D.G. (2018) Clustal Omega for making accurate alignments of many protein sequences. Protein Science, 27, 135–145.

24. Ye, K., Lameijer, E.-W.M., Beukers, M.W. and IJzerman, A.P. (2006) A two-entropies analysis to identify functional positions in the transmembrane region of class A G protein-coupled receptors. Proteins: Structure, Func4on, and Bioinforma4cs, 63, 1018–1030.

25. Ye, K., Vriend, G. and IJzerman, A.P. (2008) Tracing evolutionary pressure. Bioinforma4cs, 24, 908–915.

26. Sandberg, M., Eriksson, L., Jonsson, J., Sjöström, M. and Wold, S. (1998) New Chemical Descriptors Relevant for the Design of Biologically Active Peptides. A Multivariate Characterization of 87 Amino Acids. J. Med. Chem., 41, 2481–2491.

27. Abagyan, R. and Totrov, M. (1994) Biased probability Monte Carlo conformational searches and electrostatic calculations for peptides and proteins. J Mol Biol, 235, 983–1002.

28. Wong, A.I. and Rock, J.M. (2021) CRISPR Interference (CRISPRi) for Targeted Gene Silencing in Mycobacteria. Methods Mol Biol, 2314, 343–364.

29. Rock, J.M., Hopkins, F.F., Chavez, A., Diallo, M., Chase, M.R., Gerrick, E.R., Pritchard, J.R., Church, G.M., Rubin, E.J., Sassex, C.M., et al. (2017) Programmable transcriptional repression in mycobacteria using an orthogonal CRISPR interference playorm. Nat Microbiol, 2, 1–9.

30. Castañeda-García, A., Prieto, A.I., Rodríguez-Beltrán, J., Alonso, N., Cantillon, D., Costas, C., Pérez-Lago, L., Zegeye, E.D., Herranz, M., Plociński, P., et al. (2017) A non-canonical mismatch repair pathway in prokaryotes. Nat Commun, 8, 14246.

31. Arnold, F.M., Hohl, M., Remm, S., Koliwer-Brandl, H., Adenau, S., Chusri, S., Sander, P., Hilbi, H. and Seeger, M.A. (2018) A uniform cloning playorm for mycobacterial genetics and protein production. Sci Rep, 8, 9539.

32. García-Nafría, J., Watson, J.F. and Greger, I.H. (2016) IVA cloning: A single-tube universal cloning system exploiting bacterial In Vivo Assembly. Scientific Reports, 6, 27459.

33. Kuin, R.C.M., Lamers, M.H. and Westen, G.J.P. van (2025) PyVADesign: a Python-based cloning tool for one-step generation of large mutant libraries. 10.1101/2025.04.04.647202.

34. Green, M. R., S., J. Molecular Cloning: A laboratory manual 4th edn, Vol. 1 Ch. 9, 157–260 (Cold Spring Harbour Laboratory Press, 2012).

35. Nakata, N., Kai, M. and Makino, M. (2012) Mutation Analysis of Mycobacterial rpoB Genes and Rifampin Resistance Using Recombinant Mycobacterium smegmatis. An4microb Agents Chemother, 56, 2008–2013.

36. Baños-Mateos, S., van Roon, A.-M.M., Lang, U.F., Maslen, S.L., Skehel, J.M. and Lamers, M.H. (2017) High-fidelity DNA replication in Mycobacterium tuberculosis relies on a trinuclear zinc center. Nat Commun, 8, 855.

37. Maki, H., Mo, J.Y. and Sekiguchi, M. (1991) A strong mutator effect caused by an amino acid change in the alpha subunit of DNA polymerase III of Escherichia coli. Journal of Biological Chemistry, 266, 5055–5061.

38. Gunasingam, N. (2023) Mycobacterium smegmatis: Exploring its Similarities with Mycobacterium tuberculosis. Mycobacterial Diseases, 13, 1–1.

39. Sparks, I.L., Derbyshire, K.M., Jacobs, W.R. and Morita, Y.S. Mycobacterium smegmatis: The Vanguard of Mycobacterial Research. J Bacteriol, 205, e00337–22.

40. Sassex, C.M., Boyd, D.H. and Rubin, E.J. (2001) Comprehensive identification of conditionally essential genes in mycobacteria. Proc Natl Acad Sci U S A, 98, 12712–12717.

41. Floss, H.G. and Yu, T.-W. (2005) RifamycinMode of Action, Resistance, and Biosynthesis. Chem. Rev., 105, 621–632.

42. Ng, W.L. and Rego, E.H. (2024) A nucleoid-associated protein is involved in the emergence of antibiotic resistance by promoting the frequent exchange of the replicative DNA polymerase in M. smegmatis. bioRxiv, 10.1101/2023.06.12.544663.

43. Balint, E. and Unk, I. (2023) For the Bejer or for the Worse? The Effect of Manganese on the Activity of Eukaryotic DNA Polymerases. Int J Mol Sci, 25, 363.

44. Rock, J.M., Lang, U.F., Chase, M.R., Ford, C.B., Gerrick, E.R., Gawande, R., Coscolla, M., Gagneux, S., Fortune, S.M. and Lamers, M.H. (2015) DNA replication fidelity in Mycobacterium tuberculosis is mediated by an ancestral prokaryotic proofreader. Nat Genet, 47, 677–681.

45. Lancy, E.D., Lifsics, M.R., Kehres, D.G. and Maurer, R. (1989) Isolation and characterization of mutants with deletions in dnaQ, the gene for the editing subunit of DNA polymerase III in Salmonella typhimurium. J Bacteriol, 171, 5572–5580.

46. Slater, S.C., Lifsics, M.R., O’Donnell, M. and Maurer, R. (1994) holE, the gene coding for the theta subunit of DNA polymerase III of Escherichia coli: characterization of a holE mutant and comparison with a dnaQ (epsilon-subunit) mutant. Journal of Bacteriology, 176, 815–821.

47. Zulkower, V. and Rosser, S. (2020) DNA Features Viewer: a sequence annotation formaxng and ploxng library for Python. Bioinformatics, 36, 4350–4352.

48. Robert, X. and Gouet, P. (2014) Deciphering key features in protein structures with the new ENDscript server. Nucleic Acids Research, 42, W320–W324.

