## Supplementary Data for "Identification of determinants of high-fidelity DNA synthesis in *M. smegmatis* DnaE1 through *in silico* and *in vivo* approaches"

**Supplementary Table 1**. DNA substrates used for primer extension assay.

| Oligo | Sequence (5′ – 3′ direction) |
|---|---|
| **TempAmm** | ggtcgcgtcgAgctctgtGGACGAAGGACTCCCAAC |
| **TempCmm** | ggtgaggtagCgatgagtGGACGAAGGACTCCCAAC |
| **TempGmm** | cctcacctacGcatcactGGACGAAGGACTCCCAAC |
| **TempTmm** | ggacgcgacgTgcacagaGGACGAAGGACTCCCAAC |
| **Primer** | GTTGGGAGTCCTTCGTCC |

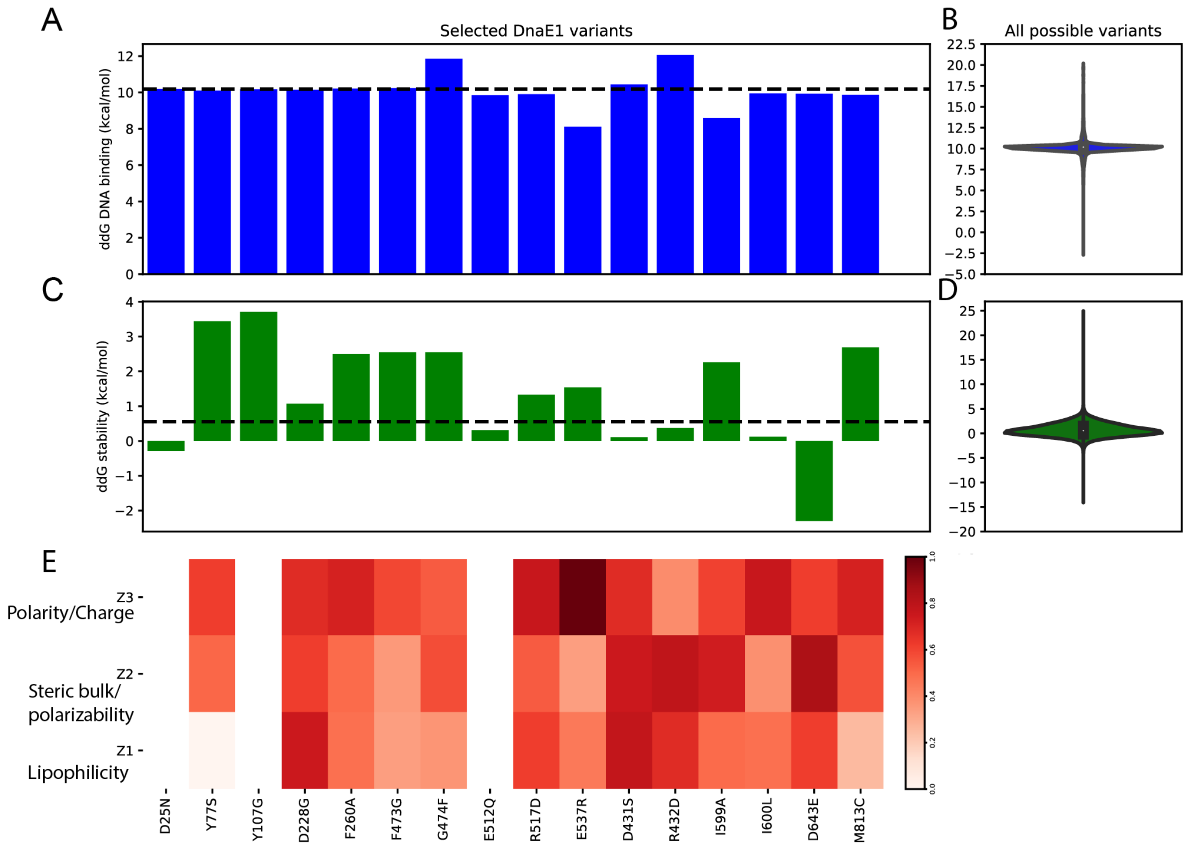


**Supplementary Figure S1**. **Properties of DnaE1 variants, based on a two-entropies analysis. (A)** Predicted change in binding free energy (ddG) for DNA binding (kcal/mol) upon mutation of a single residue for selected variants and **(B)** all possible DnaE1 variants. **(C)** Predicted effect on DnaE1 stability (ddG) upon mutation for selected variants and **(D)** all possible variants. Calculations were done in ICM-Pro version 3.9-3a (Molsoft L.L.C.) and the black dotted line represents the median value based on all possible variants. **(E)** Heatmap highlighting the differences in physicochemical properties for selected positions in the Multiple Sequence Alignment. No data is shown for control variants D25N, Y107G and for E512Q. Lipophilicity, steric properties (Steric bulk/Polarizability) and electronic properties (Polarity/Charge) are represented by Z1, Z2 and Z3 respectively (26). Scaled standard deviations of these physicochemical properties are shown, where a higher standard deviation indicates a bigger difference in the property for all residues in that position of the Multiple Sequence Alignment.

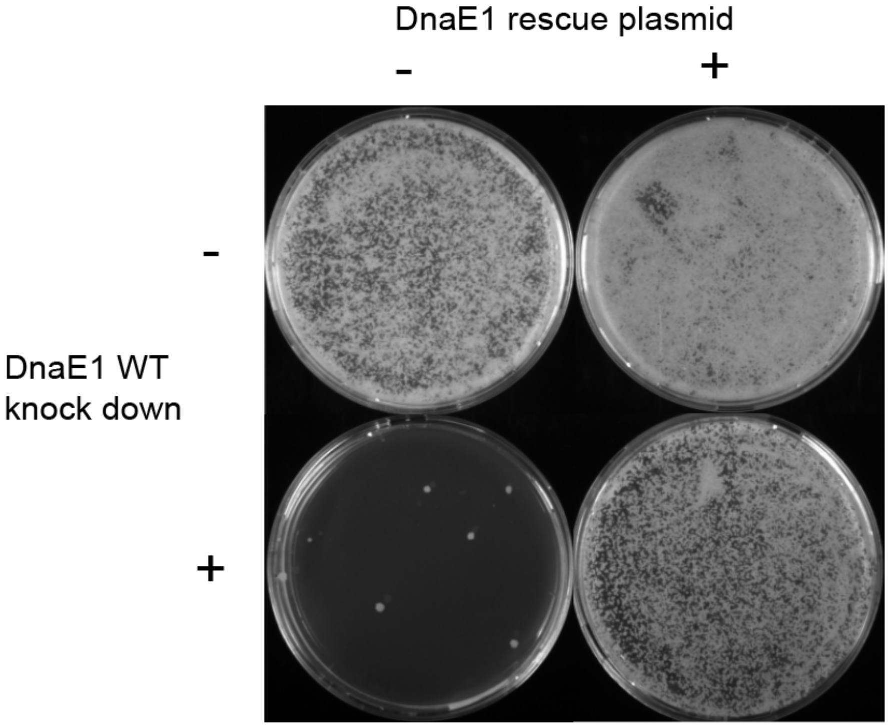


**Supplementary Figure S2. dCas9 Knock-down and rescue of DnaE1.** *M. smegmatis* cells containing dCas9 plasmid with sgRNA targeting the endogenous DnaE1 gene and a dCas9-resistant expression plasmid of DnaE1 gene in absence and presence of dCas9 and rescue DnaE1.

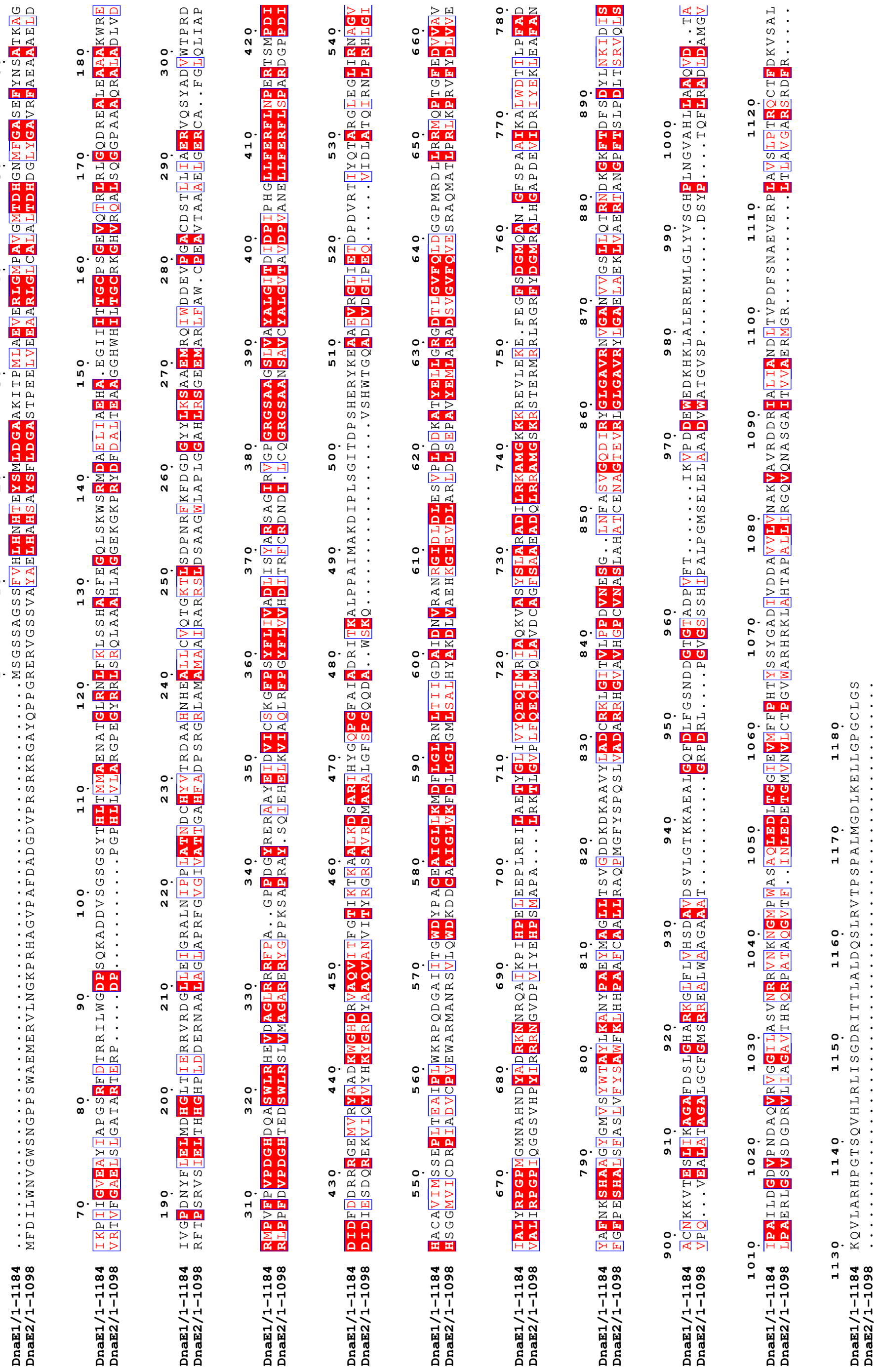

**Supplementary Figure S3. Alignment of *M. tuberculosis* DnaE1 and DnaE2 protein sequences**. Identical residues are marked in red background and white letters, similar residues are marked with red letter in blue box. Overall sequence identity = 25%, overall sequence similarity = 45%. Figure was created with ESPript version 3.0 (48).

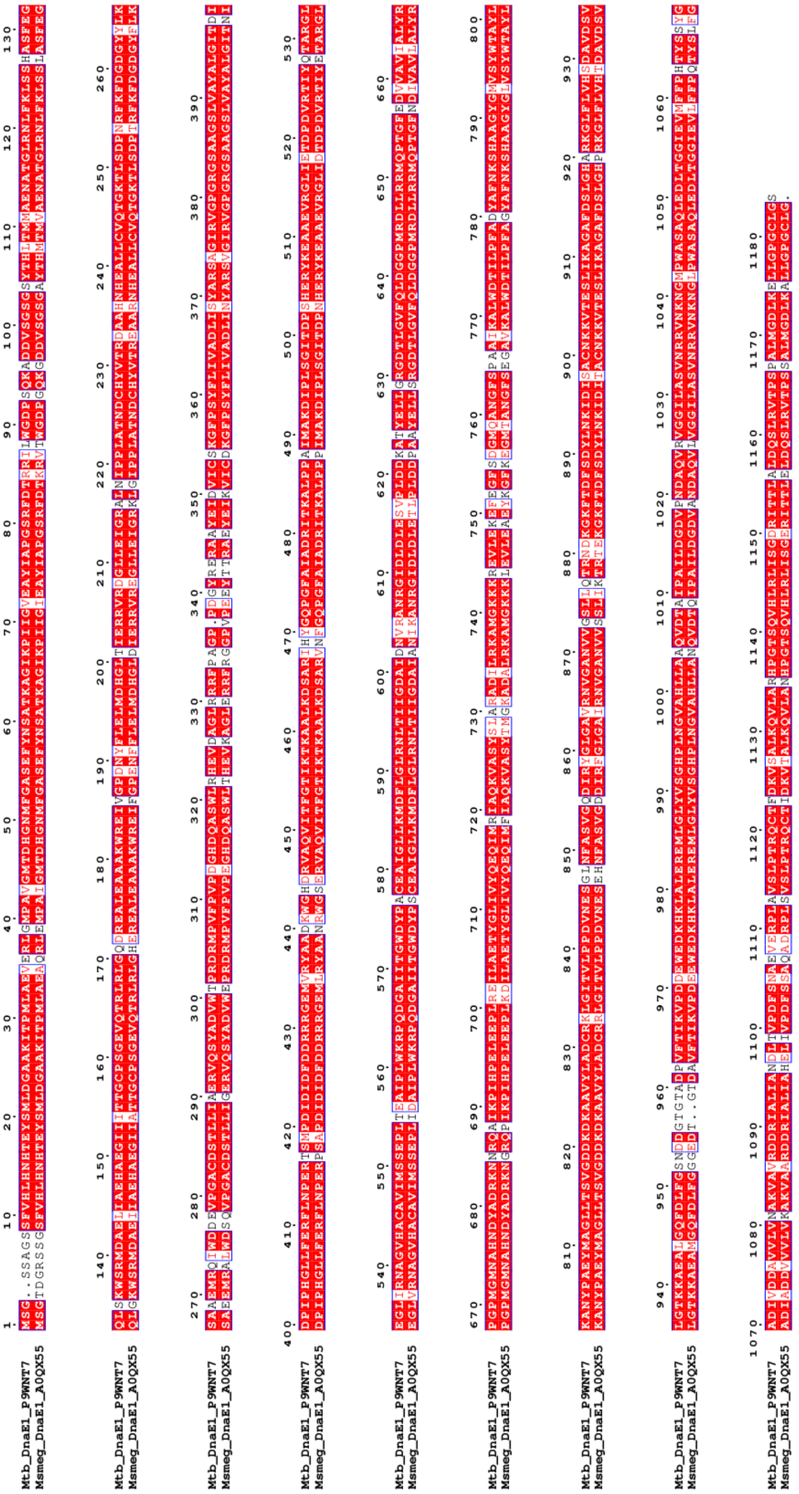

**Supplementary Figure S4.** Alignment of *M. tuberculosis* DnaE1 and *M. smegmatis* DnaE1 protein sequences. Identical residues are marked in red background and white letters, similar residues are marked with red letter in blue box. Overall sequence identity = 86%, overall sequence similarity = 96%. Figure was created with ESPript version 3.0 (48).

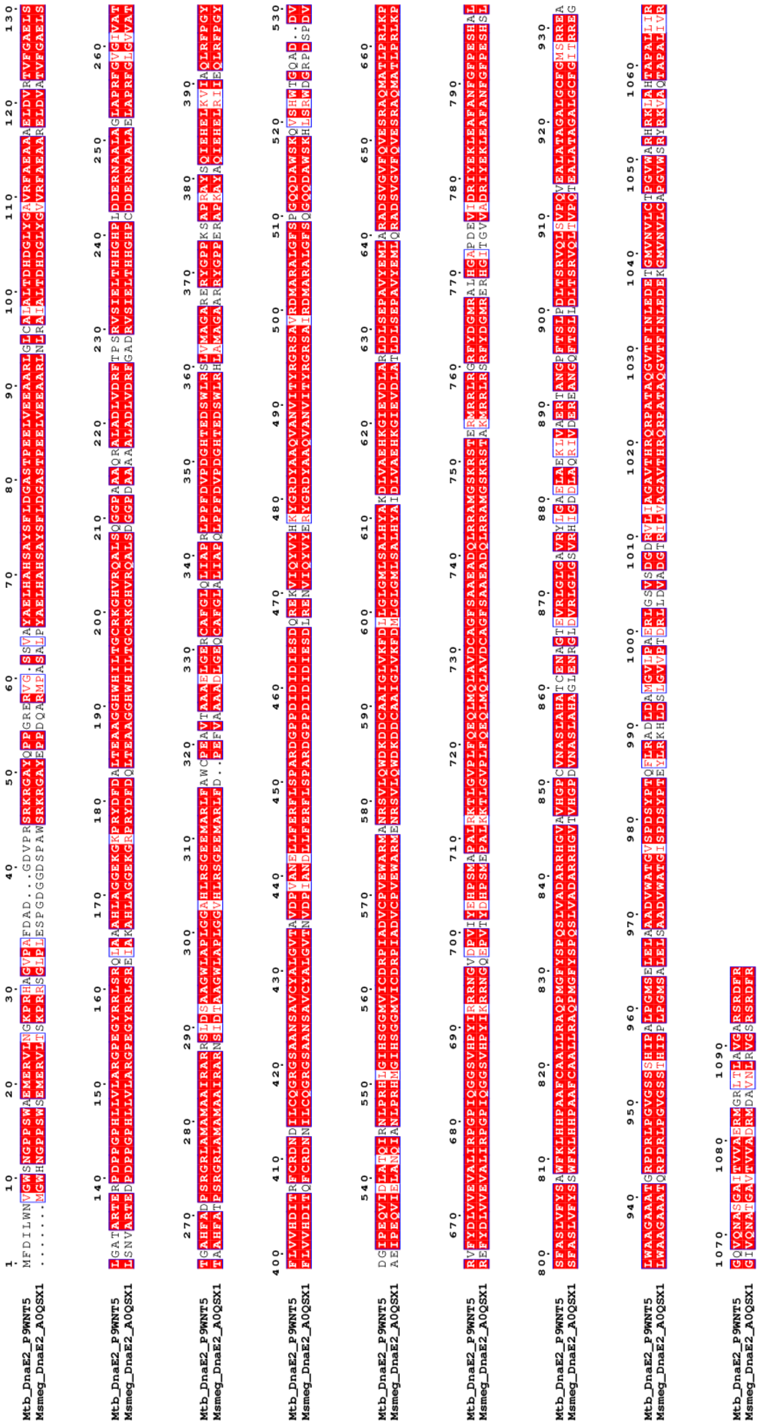

**Supplementary Figure S5.** Alignment of *M. tuberculosis* DnaE2 and *M. smegmatis* DnaE2 protein sequences. Identical residues are marked in red background and white letters, similar residues are marked with red letter in blue box. Overall sequence identity = 82%, overall sequence similarity = 94%. Figure was created with ESPript version 3.0 (48).
